# Online μSEC^2^-nRPLC-MS for improved sensitivity of intact protein detection of IEF separated non-human primate cerebrospinal fluid proteins

**DOI:** 10.1101/2021.06.08.447575

**Authors:** Erika N. Cline, Carina Alvarez, Jiana Duan, Steven M. Patrie

## Abstract

Proteoform-resolved information, obtained by top-down (TD) “intact protein” proteomics, is expected to contribute substantially to the understanding of molecular pathogenic mechanisms and in turn, identify novel therapeutic and diagnostic targets. However, the robustness of mass spectrometry (MS) analysis of intact proteins in complex biological samples is hindered by high dynamic range in protein concentration and mass, protein instability, and buffer complexity. Here, we describe an evolutionary step for intact protein investigations through the online implementation of tandem microflow size exclusion chromatography with nanoflow reversed-phase liquid chromatography and MS (μSEC^2^-nRPLC-MS). Online serial high-/low-pass SEC filtration overcomes the aforementioned hurdles to intact proteomic analysis through automated sample desalting/cleanup and enrichment of target mass ranges (5-155 kDa) prior to nRPLC-MS. The coupling of μSEC to nRPLC is achieved through a novel injection volume control (IVC) strategy of inserting protein trap columns pre- and post-μSEC columns to enable injection of dilute samples in high volumes without loss of sensitivity or resolution. Critical characteristics of the approach are tested via rigorous investigations on samples of varied complexity and chemical background. Application of the platform to cerebrospinal fluid (CSF) pre-fractionated by OFFGEL isoelectric focusing drastically increases the number of intact mass tags (IMTs) detected within the target mass range (5-30 kDa) in comparison to one-dimensional nRPLC-MS with approximately 100x less CSF than previous OFFGEL studies. Furthermore, the modular design of the μSEC^2^-nRPLC-MS platform is robust and promises significant flexibility for large-scale TDMS analysis of diverse samples either directly or in concert with other multidimensional fractionation steps.

## INTRODUCTION

Top-down mass spectrometry (TDMS) on intact proteins allows direct detection of proteoforms, i.e., the different molecular forms of a protein from a single gene, which may include combinations of post-translational modifications (PTMs), polymorphisms, homologous genes, and alternative splicing events.^1^ Different proteoforms from the same gene may participate in unique molecular interactions,^2, 3^ which suggests proteoform-level insights will substantially contribute to our understanding of the molecular mechanisms that underlie various diseases and consequentially, enable precision medicine through the identification of novel therapeutic targets and diagnostic biomarkers, particularly from biofluids such as cerebrospinal fluid (CSF).^4, 5^ Despite its potential, intact protein analysis faces many unique challenges including complexity of extraction, enrichment, purification, and separations of both small and larger proteins with diverse physiochemical properties and solubilities.^6, 7^ CSF is an ideal source for biomarkers for diseases of the central nervous system (CNS)^8, 9^ as it circulates within the ventricles of the brain, surrounds the spinal cord, and directly interchanges with the extracellular/interstitial fluid of the CNS.^10^ However, CSF biomarker discovery is challenged by the low concentration of CSF proteins (as low as 0.15 mg/mL),^11^ with CNS-specific proteins overshadowed by serum-derived proteins comprising ~40-80% (i.e., 0.06-0.12 mg/mL) of the CSF proteome.^12, 13^ In addition, the high salt concentration of CSF can interfere with various analytical techniques without sample cleanup.^12^ While sample preprocessing (e.g., molecular weight cutoff filters, dialysis) is often used for desalting^14, 15^ and affinity chromatography is often used to deplete highly abundant serum proteins,^16, 17^ these methods can be tedious and costly, with low robustness, and are prone to precipitation or sample loss at low volumes.^14, 15^

Size-based fractionation methods may offer a solution to all of these issues, i.e., removal of abundant, high molecular weight (HMW) serum proteins, such as albumin;^12, 13^ removal of low MW (LMW) salts; and general reduction of sample complexity. For TD investigations, gel-based size fractionation methods have been used, such as GELFrEE (gel elution liquid fraction entrapment electrophoresis)^18^ and PEPPI-MS (passively eluting proteins from polyacrylamide gels as intact species for MS),^19^ however these methods utilize MS-incompatible detergents (e.g., SDS), necessitating additional cleanup prior to MS. Size exclusion chromatography (SEC), on the other hand, does not exhibit the disadvantages of chromatographic preprocessing methods^14^ and can be used for removal of salts and detergents.^20–24^ Notably, offline SEC can fractionate complex biological samples prior to RPLC-TDMS analysis, achieving a significant increase in protein/proteoform detection over RPLC-TDMS alone by not only reducing the sample complexity but also decreasing ion suppression of HMW proteins when present in the mass spectrometer with LMW proteins.^25, 26^ Online SEC-MS has been achieved,^27–35^ but its widespread use, especially high resolution separations, has been precluded by a number of technical constraints. First, there are large flow rate incompatibilities between SEC (typically 150-500 μL/min for commonly utilized column diameters of 4.6-9.4 mm)^21, 24^ and sensitive downstream methods, e.g., nanoflow reversed-phase liquid chromatography (nRPLC) (0.1-1.0 μL/min) coupled with electrospray ionization (ESI)-MS. The SEC/nLC flow rate mismatch has been overcome by post-SEC flow-spitting^29^ but at a cost of high sample loss. Alternatively, this hurdle can be overcome, in part, by decreasing SEC column diameter; however, this severely limits the sample injection volume for optimal resolution due to band broadening (5-10 % of total column volume),^36, 37^ necessitating sample preprocessing to concentrate samples.

In the current report, we introduce a multidimensional MS workflow for intact proteins that couples microflow SEC (μSEC) online with nRPLC and MS. The fully integrated approach, termed μSEC^2^-nRPLC-MS, employs high- and low-pass SEC filtration with a unique automated injection volume control (IVC) technique to overcome limitations associated with integration of SEC online with sensitive nRPLC in clinical intact proteomics investigations (e.g., system pressure limitations, LMW sample buffer complexity, and broad dynamic range of protein concentration and mass). The multidimensional workflow fully automates sample desalting/cleanup and enrichment of the proteins < 30 kDa prior to nRPLC-MS analysis, a MW range that comprises many non abundant proteins in CSF^12^ and putative CSF biomarkers for various neurological disorders (e.g., proteoforms of amyloid beta, hemoglobin subunits, transthyretin, cystatin C, lipocalin prostaglandin d-synthase (L-PGDS), and myelin basic protein).^38–42^ Characteristics of the individual components and the entire platform were tested via rigorous investigations on samples of varied complexity and chemical background including CSF pre-fractionated by OFFGEL isoelectric focusing (IEF). In comparison with one dimensional (1D) reversed-phase liquid chromatography (RPLC), multi-dimensional analysis is a powerful and commonly used strategy in both top-down and bottom-up proteomic settings to improve dynamic range for detection of peptides and proteins with diverse physiochemical properties,^5–7, 43, 44^ including, two-dimensional OFFGEL IEF-nRPLC-MS which has been shown to improve protein/proteoform detection by up to 10× over 1D nRPLC-MS.^45^ IEF fractionation is especially useful for resolution of heterogenous charged modifications, e.g., notably enabling detection of > 200 glycoproteoforms of the L-type prostaglandin D-synthase in a previous study.^39^ Herein, we show that removal of IEF buffers and HMW serum proteins from CSF via two stage SEC filtration permits hundreds of intact mass tags (IMTs) to be observed from 100 μL of CSF (est. 15 μg protein), which is approximately 100x less CSF than used in previous studies.^39, 45^ Furthermore, the modular design of the μSEC^2^-nRPLC-MS platform lends robustness, flexibility, and implementation via standard chromatography equipment. This combined with the results from a variety of SEC column resins and architectures highlights the promise of the approach in large-scale TDMS analysis of diverse samples either directly or in concert with other multidimensional fractionation steps.

## MATERIALS & METHODS

### Materials

IgGs utilized were SILu (Sigma MSQC4) and anti-BNP (ICL RBNP-65A-Z). All other proteins were purchased from Sigma Aldrich: bovine serum albumin (BSA; A7906), human haptoglobin (H3536), bovine carbonic anhydrase (C2624), bovine myoglobin (M5696), bovine α-lactalbumin (L5385), and bovine ubiquitin (U6253). Peptides utilized were a custom peptide (H-GLFYVDFLSQDKVSIALSSHWINPR-OH, 2894.29 Da; Global Peptide), fibrinopeptide B (1570.6 Da; Sigma F3261), and Z-Leu-Leu-Leu-al (475.62 Da; Sigma C2211). Detergents were purchased from Sigma Aldrich: CHAPS (3-[(3-Cholamidopropyl)dimethylammonio]-1-propanesulfonate hydrate; C3023), and Triton X-100 (X100). Chromatography solvents were purchased from Fisher: water (Optima^™^ LC/MS grade; W64), acetonitrile (Optima^™^ LC/MS grade; A9554), and formic acid (Pierce^™^ LC/MS grade; PI28905).

### Isoelectric focusing (IEF)

OFFGEL IEF conditions generally followed that previously reported for the analysis of the CSF proteome.^45^ CSF from cynomolgus monkeys was a kind gift from AbbVie Inc. CSF (100 μL) proteins (est. 15 μg) or a mixture of haptoglobin (10 μg) and BSA (25 μg) was diluted in rehydration buffer (6.7 M urea, 1.0 M thiourea, 4.8 % v/v glycerol, 9.6 mg/mL dithiothreitol, 0.3 % v/v IPG ampholytes (Cytiva)) ± detergents, where indicated, and loaded onto a 24 cm Immobiline DryStrip gel (Cytiva) with pH range of 4-7 (haptoglobin/BSA) or 3-10 (CSF). Samples were focused on an Agilent 3100 OFFGEL Fractionator for 100 kVh at 22 °C (2 h at 200 V, 4 h linear gradient 200-1200 V, 41 h linear gradient 1200-8000 V) then voltage was held at 1000 V until sample collection. At collection, sample fraction volume was ~150 μL. Samples were flash-frozen and stored at −80 °C until analysis by SDS-PAGE/silver staining or LC-MS. SDS-PAGE was performed with 2.5 % v/v 2-mercaptoethanol (Sigma) on 4-15% Biorad Criterion TGX Stain-Free Tris-Glycine gels; silver staining via manufacturer’s protocol (Pierce 24612). Prior to LC-MS analysis, samples were denatured with 10 mM TCEP (tris(2-carboxyethyl)phosphine) for 1 h at 30 °C, centrifuged at 14,000 rcf/10 min, and then the supernatant was diluted 1:1 in 25-50 % v/v acetonitrile with 0.2 % v/v formic acid.

### μSEC^2^-nRPLC-MS

The μSEC^2^-nRPLC-MS workflow is shown in **Figure 1** and **Figure S1**. Briefly, samples are injected via a Dionex UltiMate 3000 autosampler and directed through various flow paths by three liquid pumps (Agilent 1100; Thermo Scientific Dionex UltiMate Nano/Cap System NCS-3500RS). A Waters Nanoacquity UPLC Symmetry C18 trap heated to 60 °C (Fisher 50781815) is used pre-SEC; a PepSwift trap is used pre-RP (200 μm x 5 mm, Thermo Scientific DX164558). Samples are eluted from the former with an injection of 10 μL acetonitrile and from the latter by an RP gradient. SEC columns contained PolyHYDROXYETHYL A^™^ (PolyHEA) resin and were purchased from polyLC: 500 Å/5 μm/150 x 1.0 mm (151HY0505); 1000 Å/5 μm/150×1.0 mm (151HY0510); 300 Å/2 μm/100×2.1 mm (102HY0203); 1000 Å/2 μm/100×2.1 mm (102HY0210); 1000 Å/2 μm/200×2.1 mm (202HY0210); 1500 Å/3 μm/200×2.1 mm (202HY0315). The RP column was purchased from Thermo Scientific (ProSwift RP-4H, 100 μm x 50 cm, #164921). Chromatography mobile phases are a mixture of “Solvent A” (95:5:0.2 water: acetonitrile: formic acid) and “Solvent B” (5:95:0.1 water: acetonitrile: formic acid). SEC mobile phases were 75A: 25B for direct SEC-MS detection or 90A: 10B when trap pre-concentration was utilized; flow rate was 30 μL/min. RPLC is performed with a linear gradient of 20-55 % B over 14 min at 1.0 μL/min. SEC or RP columns were interfaced to an LTQ Orbitrap XL/ETD with a custom ESI source and a 50 μm (± 3 μm) ESI tip (New Objective TT360-50-50-N-5). The mass spectrometer was tuned to the [M + 10H]^10+^ of ubiquitin by direct infusion. All samples, including standard mixtures and CSF samples, were analyzed in ion trap mode with the following parameters: maximum accumulation time of 50 ms with 10 μscans collected for normal mass range (< 2000 m/z) or 25 μscans for high mass range (> 2000 m/z; used for standard mixtures, only).

**Figure 1:**
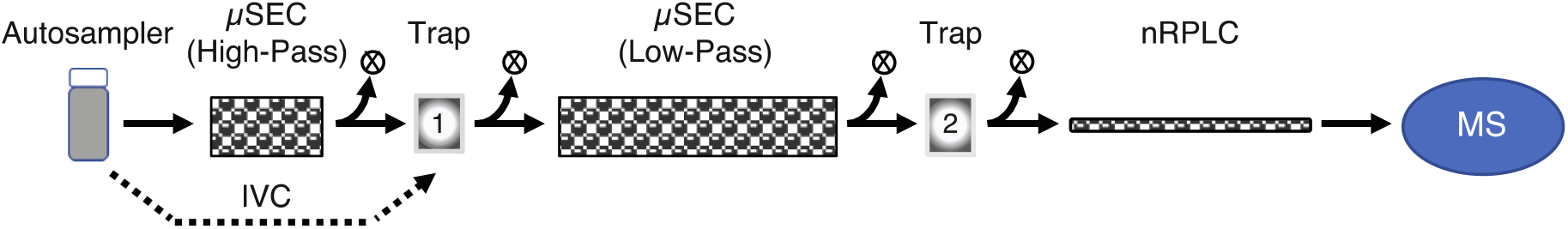
The μSEC^2^-nRPLC-MS flow path for denatured-mode proteome investigations. OFFGEL IEF fractions are injected onto the “High-Pass” (HP) column with early eluting proteins concentrated onto a RP-trap (1) and late eluting LMW buffers, salts, detergents directed to waste (X). Injection volume control (IVC; *see text*) is employed to facilitate SEC separation of LMW and HMW proteins through the “Low-Pass” (LP) column with select MW ranges concentrated onto a second RP-trap (2) for subsequent nRPLC-MS analysis. Alternate flow paths are described in **Figure S1**.

### μSEC-MS

The μSEC-MS flowpath of the platform is shown in Figure S1B-C. Protein/peptide standard mixtures typically contained SILu IgG (150 kDa), BSA (66 kDa), carbonic anhydrase (29 kDa), myoglobin (17 kDa), ubiquitin (8.6 kDa), custom peptide (2.9 kDa), fibrinopeptide B (1.6 kDa), and Z-Leu-Leu-Leu-al (0.48 kDa) and were injected in a 1 μL volume via full-loop injection mode, unless otherwise specified. Instrument parameters were identical to those stated in previous section. Extracted ion chromatograms (XIC) are plotted for individual proteins/peptides, with intensity normalized to each individual maximum and retention time (Rt) normalized to Rt_BSA_ = 1.5 min. Calibration curves plot Rt_max_ ± half-width half-maximum; third order polynomials are fit to the data.^46^

### Data Analysis

Chromatograms and mass spectra were visualized in the Thermo Xcalibur Qual Browser. Chromatography traces were plotted as total ion counts (TIC) or XIC, where indicated. Rt maxima of chromatograms were calculated automatically by Xcalibur; full width half maxima (FWHM) were measured manually in Xcalibur. Protein Deconvolution 4.0 (PD4, Thermo Fisher Scientific) was used for standard protein quantification (sum intensity output from PD4) and determination of average mass of unknown CSF proteins (i.e., intact mass tags or IMTs) within .raw files. Typical deconvolution settings were noise rejection, 99 % confidence; relative abundance threshold, 10 %; and mass tolerance, 100 ppm. CSF data were analyzed in 10 kDa target mass range increments, adjusting the following: the target mass to the median of the range; the charge state range according to the target mass range; and the minimum number of adjacent charges detected according to 50 % of the theoretical maximum charges possible within the m/z scan range (typically 700-2000 m/z) and the target mass range. The sliding window method was employed within PD4 with the following settings: target average spectrum width, 0.5 min; target average spectrum offset, 50 %; merge tolerance, 75 ppm; maximum retention time gap, 1.0 min; minimum number of detected intervals, 3. Protein identifications were made by ProSight Lite on in-source dissociation (ISD) data (source voltage = 60) obtained from a separate μSEC^2^-nRPLC-MS run, as previously outlined.^45^ P-scores were calculated as before.^47^ L-PGDS glycoproteoform assignments were made via accurate mass tags. Target cynomolgus monkey sequences were obtained from Uniprot.^48^ This included search by ISD data. Plots were prepared in Excel (v 16.45.21011103, Microsoft), Prism (v 9.0.0, GraphPad) or Xcalibur Qual Browser (v 3.1.66.10, Thermo Scientific) and figures in Illustrator (2021, Adobe). Mass spectral signal-to-noise (S/N) ratios were calculated in mMass (v5.5.0, www.mmass.org).^49^

## RESULTS AND DISCUSSION

### Performance of microflow SEC (μSEC) columns

Previously, SEC has been utilized in connection with studies on intact proteins for both online sample cleanup^28, 50^ and offline fractionation prior to RPLC-TDMS.^24–26^ Traditionally such investigations utilize analytical column diameters (e.g., ⵁ = 4.6 or 9.4 mm) at flowrates (150-500 μL/min)^21, 24^ that do not readily conform to pressure limits of downstream narrow bore RP-traps/columns without use of supplemental flow splitting techniques^29^ that can lead to significant sample loss. We postulated that for our workflow (**Figure 1**), the use of SEC in the microflow (1-100 μL/min) regime (i.e., μSEC) would overcome this flow mismatch and thus permit online integration of SEC with sensitive nanoflow RPLC (nRPLC). The two stages of μSEC are intended to help address other limitations associated with nRPLC on intact proteins. The first stage, referred to as “high-pass” (HP)-μSEC, is to help isolate proteins from background LMW buffers, salts, detergents, reducing agents, etc., in samples (such as OFFGEL IEF fractions) that readily clog or overwhelm the binding capacity of capillary traps and nRPLC columns.^51^ The second stage, supported by a novel injection volume control (IVC) strategy (*vide infra*), size separates proteins prior to subsequent MS or LC-MS analysis. For OFFGEL investigations on CSF, e.g., this stage, referred to as “low-pass” (LP)-μSEC, is intended to exclude abundant HMW serum proteins (e.g., albumin) that deleteriously impact detection of non-abundant CSF proteins/proteoforms.^12^

To examine the efficacy of the μSEC^2^-nRPLC-MS platform for denatured mode analysis, we first exploited a simple μSEC-MS detection scheme (**Figure S1B**) to examine column properties for system-level compatibility with downstream nRPLC-MS. Preliminary investigations tested PolyHYDROXYETHYL A^™^ (PolyHEA) columns of varied diameter (ⵁ = 0.5-2.1 mm), particle size (2-5 μm), and flowrates (*not shown*) with ⵁ = 2.1 mm, 2 μm particle size, and 30 μL/min flowrate selected for further investigation into other column properties most suitable for the HP or LP modes (i.e., *ℓ* = 100 vs. 200 mm; pore size = 300, 1000, or 1500 Å). The choice of PolyHEA resins was motived in part by past work that showed offline serial SEC (sSEC) successfully fractionated large sarcomere proteins prior to RPLC-TDMS.^24–26^ PolyHEA resin is a neutral, polar resin that can function for either hydrophilic interaction chromatography (HILIC) (mobile phase > 60 % organic) or native- or denatured-state SEC (mobile phase < 60 % organic), over broad mass ranges dependent upon pore _size._24-26, 52

For the various column properties tested, calibration curves fit with a third-order polynomial^46^ were generated from mixtures of standard proteins (9-150 kDa) and peptides (0.5-3 kDa) under different mobile phase conditions amenable to the starting equilibration conditions necessary for RPLC-MS of intact proteins (**Figure 2** and **Figures S2-3**). Various chromatographic parameters were assessed, including FWHM, elution time window, and resolution of albumin (66 kDa) from proteins < 30 kDa (**Tables S1-3**). To summarize, the results obtained on various column and resin architectures typically show dual linearity ranges dependent on pore size, which aligns well with theory for this resin under denaturing conditions^52^ (**Figure S2**). For example, a 1000 Å pore, *ℓ* = 200 mm column shows chromatographic peaks with an average FWHM of 12.5 ± 1.5 (SEM) sec from 0.5-66 kDa (**Figure 2A**), achieving half-maxima resolution for all proteins 9-30 kDa and baseline resolution of albumin from proteins < 30 kDa (0.24 min; **Figure 2B**). While the elution time for the smallest peptide was 11 minutes, all proteins > 3 kDa eluted within the first 5 minutes (**Figure 2A**) suggesting this column at this flow rate (30 μL/min) may have minimal impact on the duty cycle for the total proteomics platform while still providing good separation for proteins < 70 kDa. Also, reduction of *ℓ* by half (*ℓ* = 200 vs. 100 mm) decreased the total elution window by approximately half while maintaining chromatographic resolution of the proteins, again as expected from SEC theory (**Figure 2B**).^37^ Furthermore, comparisons of other pore sizes to the 1000 Å resin showed that 300 Å resin improved the separation of peptides < 3 kDa at the expense of the proteins > 3 kDa, the latter of which largely eluted together within the column void volume (< 1.5 min) (**Figure 2C**). By contrast, 1500 Å resin provided good separation between albumin (66 kDa) and an antibody (150 kDa) at the expense of resolution for proteins and peptides < 66 kDa (**Figure S2**). Notably, a subsequent online sSEC-MS investigation that linked 1000 Å resin and 300 Å resin columns (*ℓ* = 100 mm ea.) in series combined the high resolution within the 9-66 kDa range of the 1000 Å resin with that in 0.5-9 kDa range of the 300 Å resin (**Figure 2C**), although the results did not significantly differ for the single 1000 Å column at 200 mm (**Figure 2B**).

**Figure 2:**
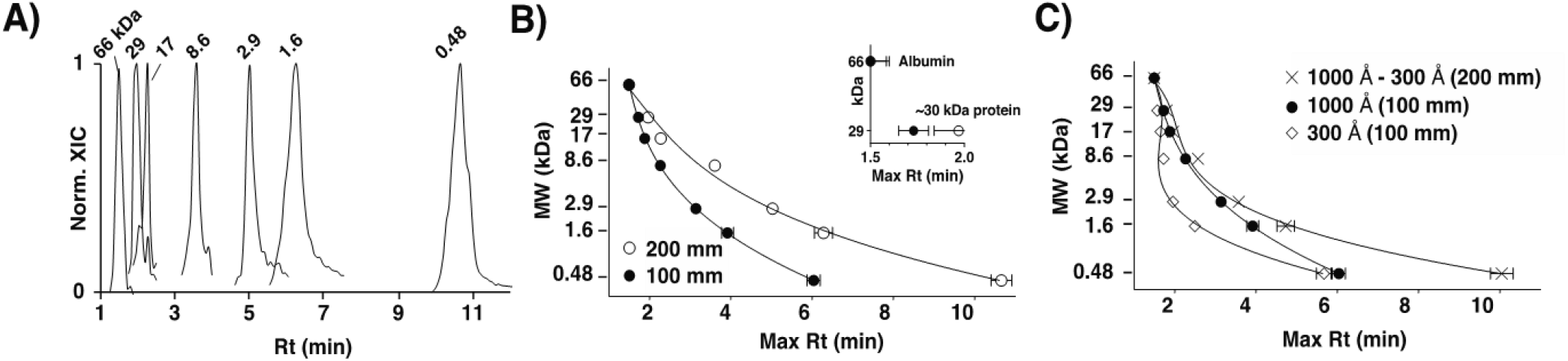
SEC performance of different PolyHEA columns. **A)** SEC-MS normalized extracted ion chromatogram (norm. XIC) for standard proteins/peptides (see Methods: μSEC-MS) run on a PolyHEA column (ⵁ = 2.1 mm, 1000 Å pore size, *ℓ* = 200 mm, 30 μL/min flowrate). **B)** SEC calibration curves compare the *ℓ* = 200 mm and *ℓ* = 100 mm columns with the separation of albumin (66 kDa) from a ~30 kDa protein (carbonic anhydrase, 29 kDa) highlighted (inset). **C)** SEC calibration curves for 300 Å, 1000 Å, and combined 1000 Å - 300 Å pore size columns. Data for other column and resin architectures are provided in the Supporting Information (**S. Tables 1-2, Figure 2**).

Overall, these investigations, supported by past investigations and SEC theory, affirm the potential of different PolyHEA resin and column architectures for denatured-mode applications.^24–26, 52^ Here, the 300 Å column (*ℓ* = 100 mm) was chosen for the first rapid HP-μSEC filtration because it effectively discriminated LMW molecules (< 3 kDa) from larger proteins (> 8 kDa) in < 2 min (**Figure 2C**). Similarly, the 1000 Å column (*ℓ* = 200 mm) was chosen for the second LP-μSEC column as it exhibited the greatest separation of albumin from proteins < 30 kDa (**Figure S2**), a common working range for large label-free quantitation investigations by TDMS.^53–56^

### Injection volume control (IVC) enables online high-resolution SEC of dilute samples while maintaining SEC-MS sensitivity

A critical barrier to the online coupling of SEC with nRPLC-MS is the small sample volume needed to achieve optimal SEC resolution—typically, 5-10% of SEC column volume (e.g., 17-35 μL for a ⵁ = 2.1, *ℓ* = 100 mm column)^36, 37^—which often necessitates use of non-ideal preparative steps to concentrate samples at the risk of poor sample recovery or use of large diameter columns (e.g., ⵁ = 4.6-9.2 mm) at high flow rates incompatible with nRPLC. We hypothesized that proteins injected in volumes greater than the volume-resolution limit could still be separated optimally by implementation of an automated IVC method that first concentrates proteins on a RP-trap and then elutes them isocratically onto a SEC column using a volume of organic solvent below the volume-resolution limit. Although online protein concentration and desalting is commonly utilized for RPLC-MS workflows in the field of proteomics and has been integrated after SEC columns to enrich targets prior to subsequent LC-MS,^28, 50^ to our knowledge, an online trap has never been utilized pre-SEC to overcome μSEC injection volume limitations.

To test this hypothesis, the impact of sample volume and concentration on SEC resolution was examined with and without pre-SEC RP-trap concentration. First, a fixed mole amount of a protein/peptide mixture was injected at a volume below (1 μL) or above (50 μL) the theoretical volume limit for high-resolution SEC. The results clearly indicate that below the volume limit high-resolution SEC is obtained (**Figure 3A**) but not above, where breakdown of the chromatographic profile for all components is observed (**Figure 3B**). Strikingly, if 50 μL of sample was first concentrated onto a RP-trap and then eluted onto the μSEC column with an injection of 100 % acetonitrile below the volume limit (10 μL), SEC resolution is restored (**Figure 3C**) for all but the most hydrophilic sample component, fibrinopeptide B (1.6 kDa), which did not bind to the trap with the current mobile phase (est. 14 % organic). Organic (acetonitrile, isopropanol) percentages of 66-100 % performed equally well for the RP-trap elution with no sample precipitation or induction of HILIC-type interactions.

**Figure 3:**
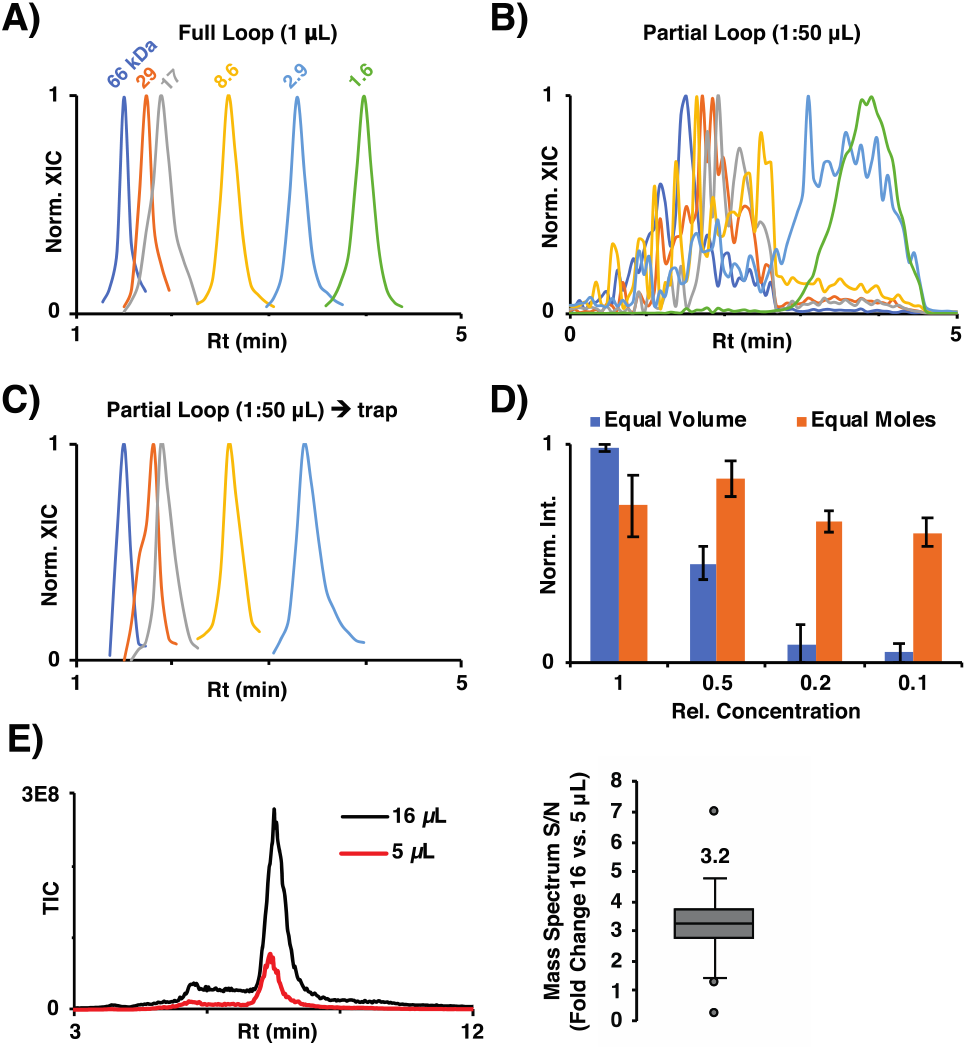
SEC IVC recovers SEC resolution and sensitivity for high volume injections. MS XIC for a six protein/peptide standard run on the *ℓ* = 100 mm, 1000 Å PolyHEA column (**Figure 2B**) w/ either 1 μL **(A)** or 50 μL **(B)** injection volumes without IVC, or w/ 50 μL injection volumes and IVC **(C)**. Description of flow paths w/ or w/o IVC are provided in the supplemental materials (**Figure S1B**). **D**) Chart compares the average summed intensity for IVC investigations on a dilution series of a four-protein standard at either constant injection volume (*blue*; 5 μL) or constant amount of each protein* (*orange*). Data is normalized to the “1x” mixture concentration ± SEM; n = 3 technical replicates. *0.80 pmol BSA, 1.14 pmol carbonic anhydrase, 1.40 pmol α-lactalbumin, and 0.18 pmol ubiquitin. **E-Left)** LC-MS TIC chromatograms for a representative CSF IEF fraction (pl = 3.3) injected at 5 or 16 μL. **Right)** Box and whisker plot highlights the distribution of MS S/N change between IMTs observed at 16 vs. 5 μL injection volume (n = 31 IMTs). The average fold improvement (3.2x) for the higher injection volume matches that expected (16/5 = 3.2).

To assess the quantitative fidelity of the IVC method, the impact of protein concentration on SEC-MS sensitivity was studied with serial dilutions of standards injected onto the RP-trap at either equal volumes or equal moles. At equal volume, the protein signal directly varies with concentration (**Figure 3D**). By contrast, for injections at equal moles delivered via varied injection volumes, the level of protein detection is largely maintained across the 10-fold dilution series. This dependence of injection amount upon data quality was further substantiated with investigations on a complex sample—an OFFGEL IEF fraction of NHP CSF—with data suggesting that improvements in LC-MS data S/N observed in a single IEF fraction directly correlate with amount injected, with an average 3.2-fold improvement in S/N for 31 IMTs when the injection volume is increased from 5 to 16 μL **(Figure 3E**). Taken together, these results suggest that IVC supports maintenance of SEC resolution and MS sensitivity even when the column sample volume limit is exceeded. Furthermore, IVC may even improve data quality of a multidimensional workflow by allowing high injection volumes and thus obviate the need for offline sample handling to concentrate the proteins in the samples.

### Introduction of HP-μSEC for removal of LMW background

The complexity of OFFGEL IEF buffers (e.g., glycerol, urea, thiourea, ampholytes, detergents) largely precludes its use in the nRPLC-MS environment without extensive sample cleanup, which can result in sample loss.^51^ To overcome this limitation in an online setting, the fidelity of the 300 Å pore size, *ℓ* = 100 mm column was examined for online HP-μSEC filtration (i.e., sample cleanup) of varied protein composition and buffer complexity. To achieve this, a switching valve was integrated post-SEC to direct the eluate either to waste or further into the multidimensional workflow (**Figure S1C**). Proof-of-concept experiments on simple protein systems (e.g., a commercial IgG antibody consisting of a high polymer background) provided preliminary evidence that this HP-μSEC filtration platform could readily remove these common buffer additives that obscured underlying protein signal (**Figure S4**). Similarly, OFFGEL IEF fractionation of the standard glycoprotein haptoglobin, with and without the detergents CHAPS or Triton X-100, showed that without filtration, only buffer components and/or the detergent are detectable via SEC-MS (**Figure 4A, black trace, peak 4**). However, with HP filtration, the LMW species are eliminated and the β- and *α*-chains of haptoglobin become readily apparent in both the chromatogram and mass spectrum (**Figure 4A; peaks 1-2,** respectively). These results on standard proteins motivated application of HP filtration to a more complex clinically relevant sample, CSF fractionation by IEF with or without CHAPS or Triton X-100. Results further corroborated those on standard proteins by showing that the 300 Å column effectively removed the buffers and composite detergents, resulting in an average of 219 IMTs detected in the 7.5-30 kDa range via RPLC-MS within the three IEF buffer conditions tested, a 95 % increase over those detected in the no-filter control (n = 6 IEF fractions) (**Figure 4B**). SEC has also proven very beneficial in TDMS analysis of mid-to-high MW proteins in past literature^25, 26^ and indeed, improved detection was also afforded here for proteins 30-155 kDa by adapting the HP method to exclude LMW buffer additives as well as small to mid-size CSF proteins <30 kDa (**Figure S5**). Taken together, these data demonstrate that this HP-μSEC filtration platform can effectively automate the removal of a variety of LMW sample components, including salts, buffers, detergents, or LMW proteins from IEF-fractionated CSF, resulting in increased protein detection.

**Figure 4:**
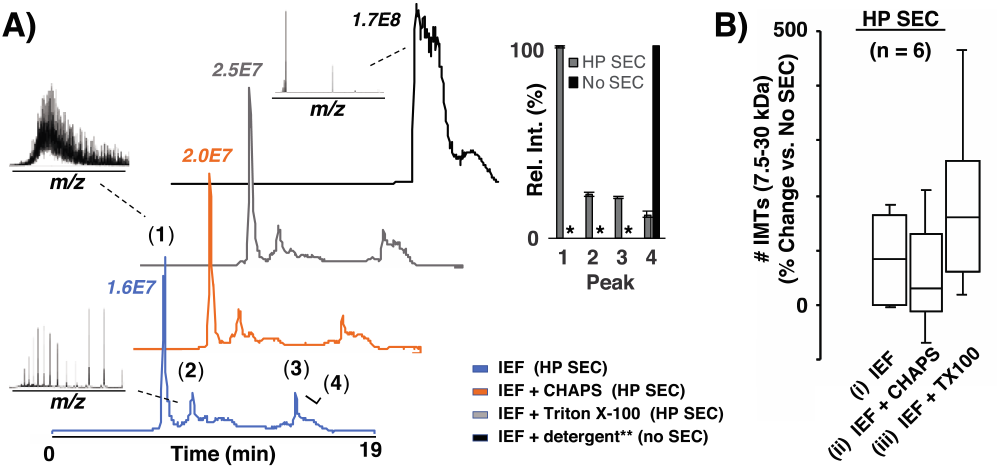
HP-μSEC filtration of IEF buffer components improves protein detection via MS. **A)** Representative μSEC-MS chromatograms w/ HP filtration on analogous IEF fractions (pI 7.6) obtained from three different OFFGEL IEF runs on the protein haptoglobin w/o detergent (*blue*) or w/ 0.07 % (v/v) CHAPS (*orange*) or 1.0 % (v/v) Triton X-100 (*gray*) added to the IEF buffer. TIC nominal values are listed above each trace Broadband mass spectrum (600-2000 *m/z*) for three observed proteins (peaks 1-3) or CHAPS detergent (peak 4). The uppermost representative detergent control chromatogram is for the same CHAPS fraction w/o high-pass (HP) filtration (*black*). A chart (inset) compares the average relative intensities of the four protein/detergent peaks w/ or w/o HP cleanup (average ± SDM; n = 3 for SEC, 1 for no SEC). *Not detected. **(B)** For an investigation on the CSF proteome, a chart compares the percent improvement in IMT detection within the 7.5-30 kDa range for IEF fractions (n = 6) w/ HP cleanup vs. w/o for three separate OFFGEL IEF runs: (i) w/o detergent or (ii) w/ CHAPS or (iii) Triton X-100 (TX100) present in the IEF run.

### μSEC^2^-nRPLC-MS platform enables dynamic range improvement via a combination of high- and low-pass filtration

Past proteomics investigations on CSF (e.g., 2D electrophoresis) have shown that removal of abundant serum proteins, many of which are mid-to-high MW (i.e., > 30 kDa) (e.g., albumin (66 kDa), *α*1-acid glycoprotein (44 kDa), *α*1-antitrypsin (45 kDa), transthyretin (55 kDa), transferrin (80 kDa), haptoglobin (86 kDa), IgGs (150 kDa), and *α*2-macroglobulin (800 kDa))^10, 12, 13, 57^ results in substantial improvement in the detection of CSF proteins that are lower in concentration and MW.^16, 17^ Furthermore, < 30 kDa is a common working range for large label-free quantitation investigations by intact protein MS.^53–56^ Therefore, we next sought to determine if the μSEC^2^-nRPLC-MS platform would impart similar benefits for CSF-IEF fractions that commonly present with these HMW proteins that can overwhelm the IEF buffering capacity at high concentration resulting in their poor fractionation.^58^ This was accomplished by integration of the 1000 Å pore size, *ℓ* = 200 mm LP-μSEC column downstream of HP-μSEC and the first RP-trap yet upstream of the second RP-trap and nRPLC-MS (**Figure 1**). The platform’s modularity, specifically ease of SEC-MS implementation, enables rapid evaluation of columns, flow conditions, and switching valve timing for loading of different SEC fractions onto traps. Thus the platform permitted comparison of three different processing schema each on seven representative IEF fractions: (1) nRPLC-MS (no SEC filtration); (2) μSEC-nRPLC-MS (HP filtration only); and (3) μSEC^2^-nRPLC-MS (combined HP-LP filtration) (**Figure S1**). In each case the mass spectra S/N and total number of IMTs above and below 30 kDa were assessed (**Figure 5** and **Figure S6A**). Chromatograms for all fractions show a predominate peak on the hydrophobic end of the RPLC gradient (**Figure S6A, top**), which typically contained HMW proteins such as albumin (66 kDa) and/or transferrin (80 kDa). With the implementation of HP or combined HP-LP filters, a progressive shift in chromatogram signal to the more polar end is observed (**Figure S6B**) that corresponds with a significant improvement in mass spectra S/N. For example, transthyretin (13 kDa; **Figure 5A-C, panel 1**)), ~20 glycoproteoforms of L-type prostaglandin D-synthase (22.4 kDa; **Figure 5A-C, panel 2**), and hemoglobin (β-chain, 15.9 kDa; **Figure 5A-C, panel 3**), identified by MS/MS analysis via ISD, had a respective 123x, 2.6x, and 22.3x-fold improvement in S/N relative to the no filtering mode. Notably, the substantial increase in S/N observed for β-hemoglobin upon HP-LP filtration corresponded with an elimination of co-eluting albumin (**Figure 5A, panel 3**). For all CSF IEF fractions, the application of HP-LP filtration vs. no filtration resulted in an average 20.5-fold decrease in mass spectrum S/N of the dominant HMW protein in each fraction along with a 35.4 % decrease in the total number of HMW IMTs (> 30 kDa; **Figure 5D-E**). In contrast, HP-LP filtration resulted in a 13.2-fold increase in mass spectrum S/N and 167 % increase in number of detected LMW IMTs (< 30 kDa; **Figure 5D-E**). These data corroborate past investigations that show the elimination of abundant proteins within samples with skewed protein dynamic range improves the observational capacity of proteomics workflows.^16, 17^ Importantly, we empirically found there was no significant loss in either HP or LP column performance after >100 CSF sample injections suggesting the PolyHEA resins are robust materials for routine intact protein separations out of complex backgrounds.

**Figure 5:**
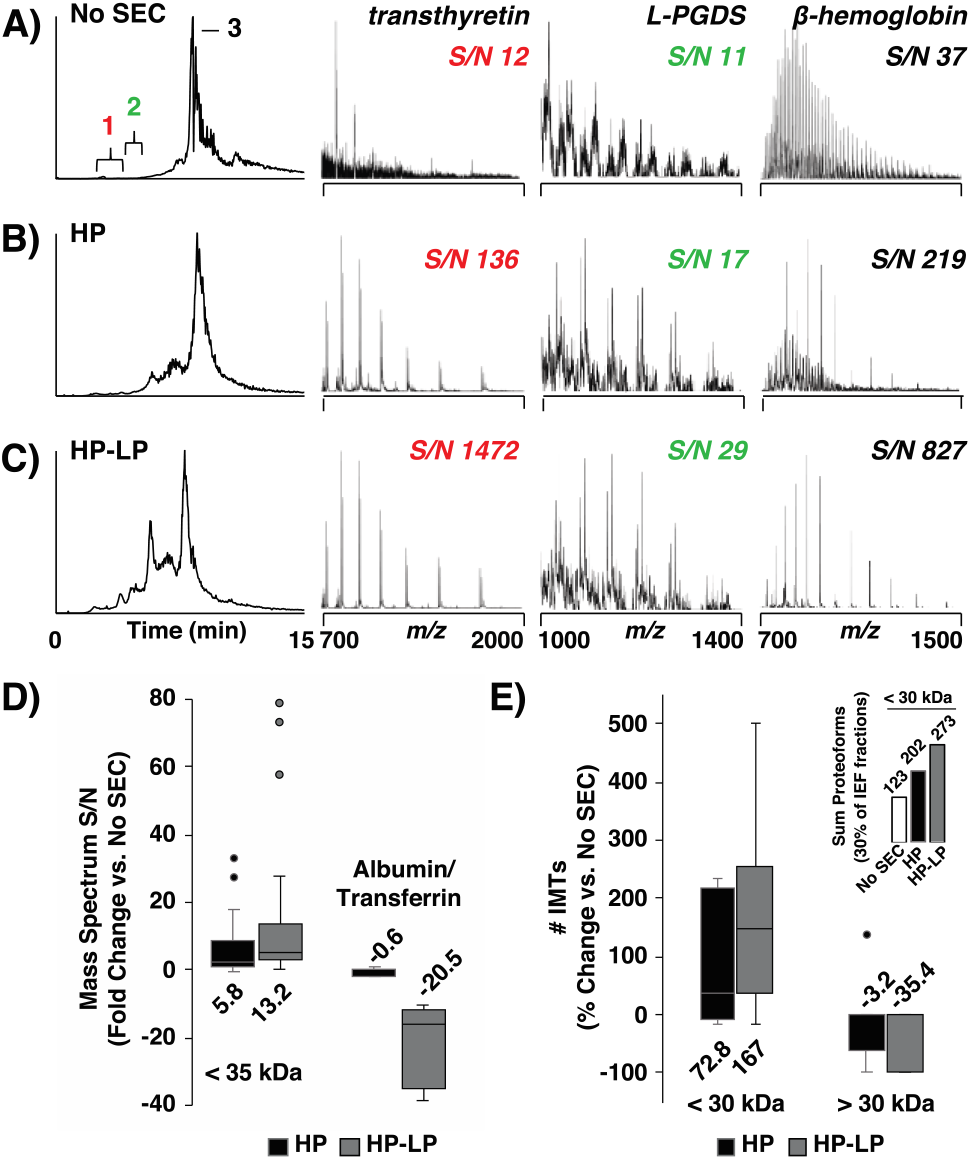
HP and LP filtration increase nRPLC-MS dynamic range for CSF IMTs < 30 kDa. LC-MS chromatograms and mass spectra at three time intervals for a representative IEF fraction (pI 8.5) analyzed **(A)** w/o SEC (“No SEC”); **(B)** w/ high-pass μSEC (“HP”); and **(C)** w/ both high- and low-pass μSEC (“HP-LP”). Proteins were identified from separate nRPLC-MS runs with ISD (**Table S4)**: transthyretin (13 kDa; **panel 1**), glycoproteoforms of L-type prostaglandin D-synthase (22.4 kDa; **panel 2**), and hemoglobin (β-chain, 15.9 kDa; **panel 3**). **D)** A box and whisker plot highlights, for the IEF fractions sampled with each mode (n = 7), the fold change in spectral S/N between the HP and HP-LP versus No SEC runs for IMTs < 35 kDa (n = 37) and the predominate HMW species observed in each chromatogram (i.e., albumin/transferrin) (**Figure S6**). **E)** The percent change in the total number of IMTs observed between the HP and HP-LP versus No SEC runs from 7.5-30 kDa and 30-70 kDa (inset; n = 7 IEF fractions).

A previous implementation of OFFGEL IEF and RPLC-MS for TDMS investigations on 1.2 mg of human CSF proteins detected ~800 proteins/proteoforms in the mass range of 5–30 kDa across 24 IEF fractions, a ~6x improvement over 1D RPLC-MS alone.^45^ Here, μSEC^2^-nRPLC-MS analysis was performed on only an estimated 15 μg (0.1 mL) of NHP CSF proteins, which resulted in detection of 273 IMTs in a similar mass range (7.5-30 kDa) from only 30 % of all the IEF fractions collected for the entire CSF proteome (est. observational capacity of 936 IMTs for entire proteome). Considering the volume of each fraction analyzed by μSEC^2^-nRPLC-MS was 36 % less than the previous report, we estimate that the inclusion of the two stage μSEC filtration effectively scaled the intact protein workflow to a nRPLC environment with a corresponding reduction of sample requirements by ~100x without loss of protein/proteoform detection. To further examine the improved sensitivity of the μSEC^2^-nRPLC-MS platform, we sought to recapitulate our previous work that characterized > 50 di-N-glycsoylated L-PGDS glycoproteoforms (> 200 proteoforms in total) from ~1.2 mg human CSF proteins^39^ using ~80-fold less protein, i.e., 15 μg (0.1 mL) of NHP (cynomolgus monkey) CSF (**Figure S7**). Here, across the pI 6-9 range of the IEF fractions 47 putative di-N-glycosylated L-PGDS glycoproteoforms were detected from ~275 putative NHP L-PGDS IMTs in total. These IMTs exhibited a similar monosaccharide composition and intensity distribution to that of human L-PGDS with the exception that NHP glycoproteforms presented with the N-glycolylneuraminic acid form of sialic acid in place of the N-acetylneuraminic acid variant which is more commonly observed in humans. Of note, because of the complexity of the glycosylation profile none of these glycoproteoforms were resolved by 1D RPLC-MS alone (data not shown). Because of the observed relationship between the amount injected on LC-MS data quality (**Figure 3E**) and the fact that the bulk of our analysis was performed with at most 10 % of each IEF fraction, prospective improvements in the detection of non-abundant proteins/proteoforms or further reduction in CSF sample requirements may be realized by expansion of the IVC approach to enable injection of higher IEF fraction volumes, which is expected to be especially beneficial for rodent studies where only ~40-150 μL CSF can be routinely collected.^59, 60^

Future directions for the μSEC^2^-nRPLC-MS workflow include adaptation to a quaternary LC system in order to allow for both high and low flow management within one LC system. Plus, future studies will aim to determine whether further sensitivity improvements may be realized by sequential μSEC^2^-nRPLC-MS on multiple MW ranges that are narrower than that employed in the current study (7.5-30 kDa), or even target specific studies, as opposed to the simple discrimination of HMW proteins from LMW performed herein. Finally, given that CSF is an ideal source for LMW biomarkers originating from the brain, the existing workflow is well positioned to examine proteoform level insights in relation for CNS disorders (e.g., differential glycosylation profiles of L-PGDS in Alzheimer’s disease, Parkinson’s disease, etc.)

## CONCLUSIONS

The unique μSEC^2^-nRPLC-MS platform provides a number of advantages for MS analysis of intact proteins within complex samples. Through automated desalting (via HP-μSEC) and concentration of the target MW range (via LP-μSEC) for the chosen clinical proteomic application, CSF pre-fractionated by OFFGEL IEF, the platform substantially increased protein/proteoform detection from sample requirements that were ~100x less than used in previous reports. The HP filtration functionality consistently improved sample analysis through the effective removal of various types of small molecules from the sample including buffers, polymers, and detergents. The utility of the LP filtration functionality was demonstrated by the removal of abundant HMW serum proteins from CSF, resulting in a drastic increase in sensitivity for detection of CSF proteins/proteoforms in the target < 30 kDa range. The robustness of the system was empirically observed with no significant loss in SEC filtration performance after > 100 CSF sample injections. Ultimately, through its modularity and flexibility in SEC column architectures, this μSEC^2^-nRPLC-MS platform is expected to be applicable to a wide variety of biological samples and target applications, serving to increase protein/proteoform detection in a high-throughput manner with minimal sample handling.

## ASSOCIATED CONTENT

### Supporting Information

The Supporting Information is available free of charge on the ACS Publications website.

μSEC-MS metrics of PolyHEA columns (Table S1); Resolution of serum albumin from proteins < 30 kDa on PolyHEA columns (Table S2); μSEC-MS metrics at different mobile phase organic content (Table S3); MS/MS identified proteins (Table S4); μSEC^2^-nRPLC-MS platform (Figure S1); Performance of different PolyHEA columns (Figure S2); Comparison of SEC mobile phase organic content (Figure S3); SEC-MS platform resolves LMW additives from proteins (Figure S4); μSEC-MS platform improves detection of HMW proteins (> 30 kDa) by elimination of LMW additives and proteins (< 30 kDa); High- and low-pass filtration increases dynamic range of CSF protein detection from OFFGEL IEF fractions (Figure S6); μSEC^2^-nRPLC-MS platform enables detection of di-N-glycosylated L-PGDS from OFFGEL IEF fractions obtained on NHP CSF (Figure S7) (PDF).

## AUTHOR INFORMATION

### Author Contributions

ENC contributed to data collection, data analysis, and writing. CA contributed to data collection. JD contributed to data collection and data analysis. SMP contributed to project concept, oversight, data analysis, and writing.

## ACKNOWLEDGMENT

The authors have declared no conflict of interests. This work was supported by the National Institute of General Medical Sciences of the National Institutes of Health under award number 1R01GM115739-01A1 and R01GM115739-04S1. Any opinions, findings, conclusions or recommendations expressed in this material are those of the authors and do not necessarily reflect the views of the National Institutes of Health. This work was also supported by the Multiple Sclerosis Society [PP-1503-04034], The Darrel K. Royal Research Fund for Alzheimer’s Disease [48680-DKR], The Texas Alzheimer’s Research and Care Consortium Investigator Grant Program [354091], and the UT System Neuroscience and Neurotechnology Research Institute [363027]. Funding was also provided by the University of Texas at Dallas, the John L. Roach Scholarship in Biomedical Research, and the Friends of Alzheimer’s Disease Research Award. The authors would like to thank Andrew J. Alpert (polyLC Inc.) for guidance in the use of PolyHEA columns for SEC and AbbVie Inc. for the kind gift of NHP CSF.

## TOC Figure

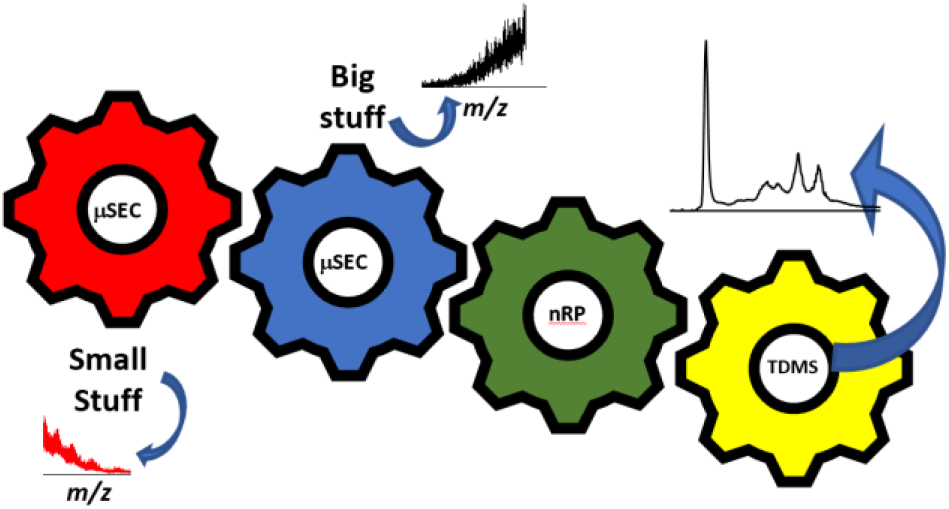

