## Supporting Information for "Online μSEC^2^-nRPLC-MS for improved sensitivity of intact protein detection of IEF separated non-human primate cerebrospinal fluid proteins"

**Table of Contents**


| | | |
|---|---|---|
| **Table S1:** | µSEC-MS metrics of PolyHEA columns | *pS3* |
| **Table S2:** | Resolution of serum albumin from proteins < 30 kDa on PolyHEA columns | *pS4* |
| **Table S3:** | µSEC-MS metrics at different mobile phase organic content | *pS4* |
| **Table S4:** | MS/MS identification of proteins highlighted in Figure 5 | *pS5-6* |
| **Figure S1:** | µSEC$^2$-nRPLC-MS platform | *pS7* |
| **Figure S2:** | Performance of different PolyHEA columns | *pS8* |
| **Figure S3:** | Comparison of SEC mobile phase organic content | *pS9* |
| **Figure S4:** | µSEC-MS platform resolves LMW additives from proteins | *pS10* |
| **Figure S5:** | µSEC-MS platform improves detection of HMW proteins (> 30 kDa) by elimination of LMW additives and proteins (< 30 kDa) | *pS11* |
| **Figure S6:** | High- and low-pass filtration increases dynamic range of CSF IMT detection from OFFGEL IEF fractions | *pS12* |
| **Figure S7:** | µSEC$^2$-nRPLC-MS platform enables detection of di-N-glycosylated L-PGDS from OFFGEL IEF fractions obtained on NHP CSF | *pS13* |
| **References** | | *pS14* |

**Supplementary Tables**

| | | 300 Å 100 mm | | 1000 Å 100 mm | | 1000 Å 200 mm | | 1500 Å 200 mm | | 1000 Å - 300 Å 100 mm each | |
|---|---|---|---|---|---|---|---|---|---|---|---|
| **Protein** | **Mass (kDa)** | **Rt (min)** | **FWHM (min)** | **Rt (min)** | **FWHM (min)** | **Rt (min)** | **FWHM (min)** | **Rt (min)** | **FWHM (min)** | **Rt (min)** | **FWHM (min)** |
| **IgG** | 150 | --- | --- | 1.00 | 0.68 | --- | --- | 0.61 | 0.42 | --- | --- |
| **Serum albumin, bovine** | 66 | 1.50 | 0.15 | 1.50 | 0.17 | 1.50 | 0.20 | 1.50 | 0.29 | 1.50 | 0.09 |
| **Carbonic anhydrase, bovine** | 29 | 1.57 | 0.15 | 1.73 | 0.16 | 1.97 | 0.26 | 1.52 | 0.14 | 1.80 | 0.16 |
| **Myoglobin, equine** | 17 | 1.65 | 0.12 | 1.88 | 0.19 | 2.28 | 0.17 | 1.84 | 0.43 | 1.96 | 0.18 |
| **Ubiquitin, bovine** | 8.6 | 1.73 | 0.14 | 2.27 | 0.15 | 3.61 | 0.20 | 2.5 | 0.43 | 2.57 | 0.19 |
| **Custom Peptide** | 2.9 | 1.96 | 0.16 | 3.14 | 0.18 | 5.03 | 0.24 | --- | --- | 3.58 | 0.22 |
| **Fibrinopeptide B** | 1.6 | 2.50 | 0.22 | 3.92 | 0.29 | 6.28 | 0.45 | 4.78 | 3.35 | 4.73 | 0.43 |
| **Z-Leu-Leu-Leu-al** | 0.48 | 5.67 | 0.36 | 6.04 | 0.32 | 10.66 | 0.51 | 6.46 | 0.74 | 10.05 | 0.56 |

**Table S1: µSEC-MS metrics of PolyHEA columns.** Maximum retention time (Rt) and peak width, measured as full-width half-maxima (FWHM), are listed for each standard protein/peptide on each column architecture tested in **Figure 2** and **Figure S2**.

| Column | Half-maxima Separation of Serum Albumin from Proteins < 30 kDa (min) |
|---|---|
| 300 Å, 100 mm | -0.08 |
| 1000 Å, 100 mm | 0.07 |
| 1000 Å, 200 mm | 0.24 |
| 1500 Å, 200 mm | -0.20 |
| 1000 Å – 300 Å, 100 mm ea. | 0.18 |

**Table S2: Resolution of serum albumin from proteins < 30 kDa on PolyHEA columns.** Separation of serum albumin (bovine; 66 kDa) from proteins < 30 kDa, as represented by a 29 kDa protein (carbonic anhydrase) is highlighted, calculated as the half-maxima separation: ($Rt_{BSA}$ – $HWHM_{BSA}$) - ($Rt_{29kDa}$ – $HWHM_{29kDa}$). A negative value indicates peak overlap (i.e., no resolution). PolyHEA column architectures are those tested in **Figure 2**, **Figure S2**, and **Table S1**.

| | | 14% ACN | | 28% ACN | |
|---|---|---|---|---|---|
| **Protein** | **Mass (kDa)** | **Rt (min)** | **FWHM (min)** | **Rt (min)** | **FWHM (min)** |
| **BSA** | 66 | 1.5 | 0.22 | 1.5 | 0.23 |
| **Carbonic anhydrase** | 29 | 1.81 | 0.16 | 1.73 | 0.18 |
| **Myoglobin** | 17 | 1.88 | 0.2 | 1.89 | 0.22 |
| **Ubiquitin** | 8.6 | 2.58 | 0.18 | 2.27 | 0.12 |
| **Peptide** | 2.9 | 3.37 | 0.22 | 3.12 | 0.17 |
| **Fibrinopeptide B** | 1.6 | 4.06 | 0.28 | 3.89 | 0.27 |
| **Z-Leu-Leu-Leu-al** | 0.48 | 6.24 | 0.38 | 6.13 | 0.35 |

**Table S3: µSEC-MS metrics at different mobile phase organic content.** Maximum retention time (Rt) and peak width, measured as full-width half-maxima (FWHM), for protein/peptide standards run on a PolyHEA column ($\varnothing$ = 2.1 mm, 1000 Å pore size, $\ell$ = 100 mm, 30 µL/min flowrate) with different mobile phase organic solvent content (14, 28 % acetonitrile, ACN); µSEC-MS data shown in **Figure S4**.

| Protein | UniProt Accession | p-score[1] | # b ions | # y ions | Ave Error (ppm) | STD Error (ppm) |
|---|---|---|---|---|---|---|
| Transthyretin | A0A0D9RZ43 | 3.30E-32 | 22 | 7 | 2.567 | 2.647 |
| Glutathione-independent PGD synthase | A0A0D9RU86 | 2.30E-13 | 4 | 5 | 0.950 | 1.813 |
| Hemoglobin subunit beta | P02028 | 9.00E-09 | 2 | 9 | 1.254 | 3.080 |
| **Transthyretin Fragment matches** | | | | | | |
| **Name** | **Ion Type** | **Ion Number** | **Theoretical Mass (Da)** | **Observed Mass (Da)** | **Mass Difference (Da)** | **Mass Difference (ppm)** |
| B9 | B | 9 | 870.408 | 870.405 | -0.003 | -3.380 |
| B10 | B | 10 | 973.417 | 973.421 | 0.003 | 3.254 |
| B12 | B | 12 | 1183.554 | 1183.555 | 0.001 | 0.986 |
| B13 | B | 13 | 1314.595 | 1314.597 | 0.002 | 1.414 |
| B16 | B | 16 | 1640.827 | 1640.827 | 0.001 | 0.316 |
| B21 | B | 21 | 2195.144 | 2195.154 | 0.01 | 4.656 |
| B22 | B | 22 | 2252.166 | 2252.183 | 0.017 | 7.626 |
| B23 | B | 23 | 2339.198 | 2339.197 | -0.001 | -0.348 |
| B25 | B | 25 | 2507.288 | 2507.310 | 0.023 | 9.042 |
| B27 | B | 27 | 2720.399 | 2720.406 | 0.007 | 2.482 |
| B28 | B | 28 | 2819.467 | 2819.474 | 0.006 | 2.211 |
| B29 | B | 29 | 2890.504 | 2890.519 | 0.015 | 5.017 |
| B30 | B | 30 | 2989.573 | 2989.576 | 0.003 | 1.166 |
| B31 | B | 31 | 3103.616 | 3103.621 | 0.005 | 1.673 |
| B32 | B | 32 | 3202.684 | 3202.706 | 0.022 | 6.745 |
| B36 | B | 36 | 3676.980 | 3676.988 | 0.008 | 2.277 |
| B39 | B | 39 | 3992.086 | 3992.080 | -0.006 | -1.499 |
| B40 | B | 40 | 4093.134 | 4093.144 | 0.011 | 2.578 |
| B42 | B | 42 | 4350.250 | 4350.280 | 0.029 | 6.747 |
| B58 | B | 58 | 5948.024 | 5948.045 | 0.021 | 3.555 |
| B63 | B | 63 | 6537.247 | 6537.256 | 0.009 | 1.396 |
| B109 | B | 109 | 11727.840 | 11727.840 | 0.008 | 0.692 |
| Y7 | Y | 7 | 785.428 | 785.429 | 0.001 | 1.410 |
| Y8 | Y | 8 | 856.465 | 856.468 | 0.003 | 3.033 |
| Y9 | Y | 9 | 957.513 | 957.516 | 0.003 | 2.654 |
| Y10 | Y | 10 | 1058.561 | 1058.563 | 0.002 | 2.014 |
| Y11 | Y | 11 | 1145.593 | 1145.597 | 0.004 | 3.441 |
| Y12 | Y | 12 | 1308.656 | 1308.658 | 0.002 | 1.177 |

| | | | | | | |
|---|---|---|---|---|---|---|
| Y15 | Y | 15 | 1655.804 | 1655.808 | 0.004 | 2.117 |
| **Glutathione-independent PGD synthase Fragment Matches** | | | | | | |
| **Name** | **Ion Type** | **Ion Number** | **Theoretical Mass** | **Observed Mass** | **Mass Difference (Da)** | **Mass Difference (ppm)** |
| B8 | B | 8 | 781.397 | 781.398 | 0.001 | 1.029 |
| B9 | B | 9 | 909.456 | 909.458 | 0.002 | 2.466 |
| B11 | B | 11 | 1120.551 | 1120.553 | 0.002 | 1.975 |
| B12 | B | 12 | 1267.620 | 1267.619 | -0.001 | -0.487 |
| Y7 | Y | 7 | 853.331 | 853.333 | 0.002 | 2.051 |
| Y8 | Y | 8 | 954.379 | 954.381 | 0.003 | 2.886 |
| Y10 | Y | 10 | 1179.490 | 1179.493 | 0.002 | 2.120 |
| Y11 | Y | 11 | 1292.574 | 1292.572 | -0.002 | -1.883 |
| Y66 | Y | 66 | 7586.636 | 7586.624 | -0.012 | -1.603 |
| **Hemoglobin subunit beta Fragment Matches** | | | | | | |
| **Name** | **Ion Type** | **Ion Number** | **Theoretical Mass** | **Observed Mass** | **Mass Difference (Da)** | **Mass Difference (ppm)** |
| B6 | B | 6 | 718.354 | 718.354 | -0.001 | -0.710 |
| B7 | B | 7 | 847.397 | 847.398 | 0.001 | 0.698 |
| Y6 | Y | 6 | 767.408 | 767.407 | -0.001 | -0.735 |
| Y7 | Y | 7 | 838.445 | 838.447 | 0.002 | 2.796 |
| Y8 | Y | 8 | 952.488 | 952.493 | 0.005 | 4.889 |
| Y9 | Y | 9 | 1023.525 | 1023.526 | 0.001 | 0.776 |
| Y10 | Y | 10 | 1122.593 | 1122.603 | 0.010 | 8.632 |
| Y11 | Y | 11 | 1179.615 | 1179.613 | -0.002 | -1.846 |
| Y12 | Y | 12 | 1250.652 | 1250.652 | -0.000 | -0.129 |
| Y20 | Y | 20 | 2138.138 | 2138.137 | -0.001 | -0.585 |
| Y23 | Y | 23 | 2462.318 | 2462.318 | 0.000 | 0.002 |

**Table S4: MS/MS identification of proteins highlighted in Figure 5.** MS/MS analysis via in-source dissociation (ISD) of representative proteins shown in Figure 5. Data is summarized for each protein at the top of the table followed by a list of each observed ion.

### Supplementary Figures

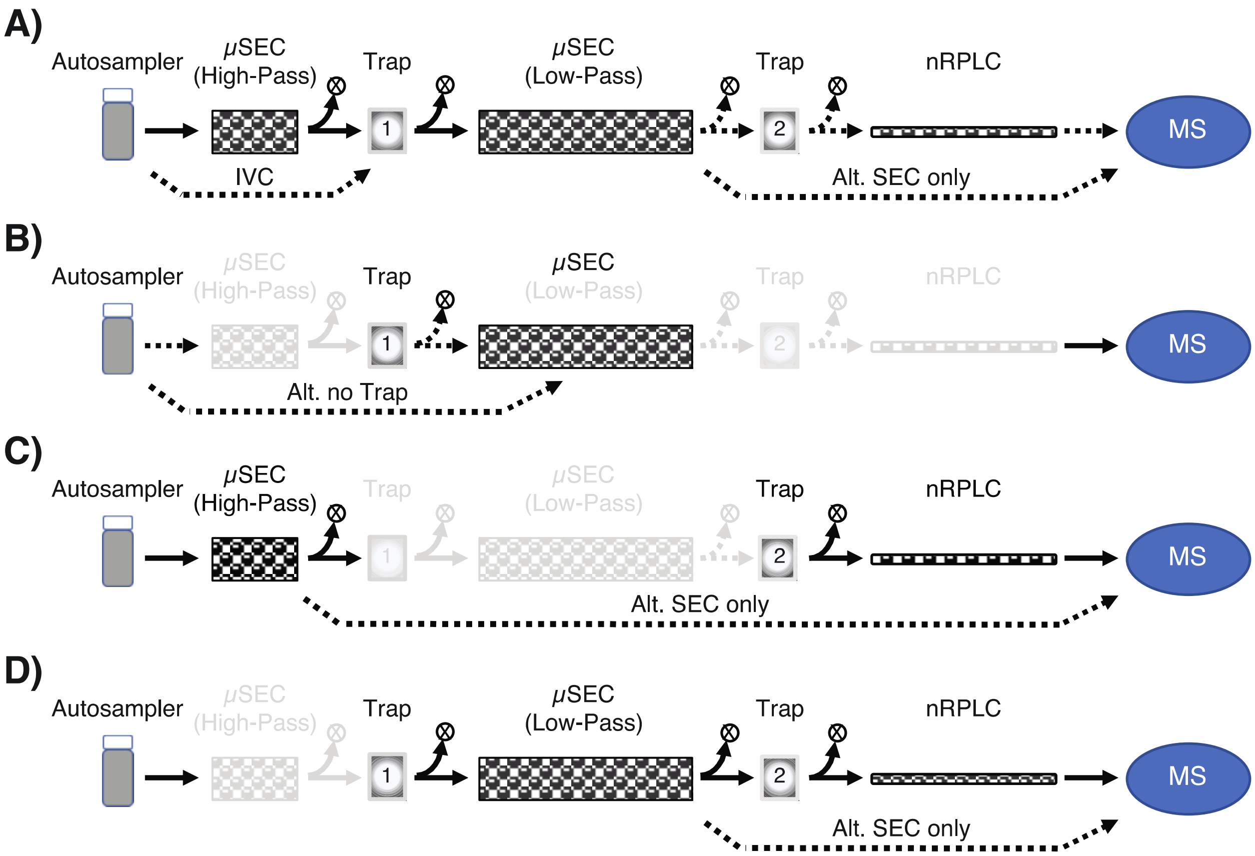


**Figure S1: μSEC$^2$-nRPLC-MS platform.** The μSEC$^2$-nRPLC-MS platform offers 8 different LC-MS detection modes. **A)** Proteome investigations typically use the complete μSEC$^2$-nRPLC-MS workflow, which includes two stages of μSEC (μSEC$^2$) for high- (HP) and low- (LP) pass filtration with a nRPLC column. During filtration, off-target fractions are sent to waste (X) while target fractions are collected onto traps. Samples are eluted by injection volume control (IVC) from Trap 1 by a subsequent injection of organic solvent (dashed arrow) and from Trap 2 by a RP gradient. The nRPLC column can be bypassed if desired, as indicated by a dashed arrow. **B-D)** In addition, integrated switching valves and a reversed-phase trap before and after the second μSEC column creates high flow path modularity permitting use of one or more columns inline with the mass spectrometer. For example, μSEC-MS can be utilized ± trap pre-concentration (i.e., IVC) for analysis of samples with various volumes, concentrations, and solvent compositions **(B)**. Alternatively, a single μSEC column, ± trap pre-concentration (i.e., IVC), can be utilized with downstream nRPLC for varying HP and LP functionalities (μSEC-nRPLC-MS) (**C-D**). Grayed out components are not used in the current configuration.

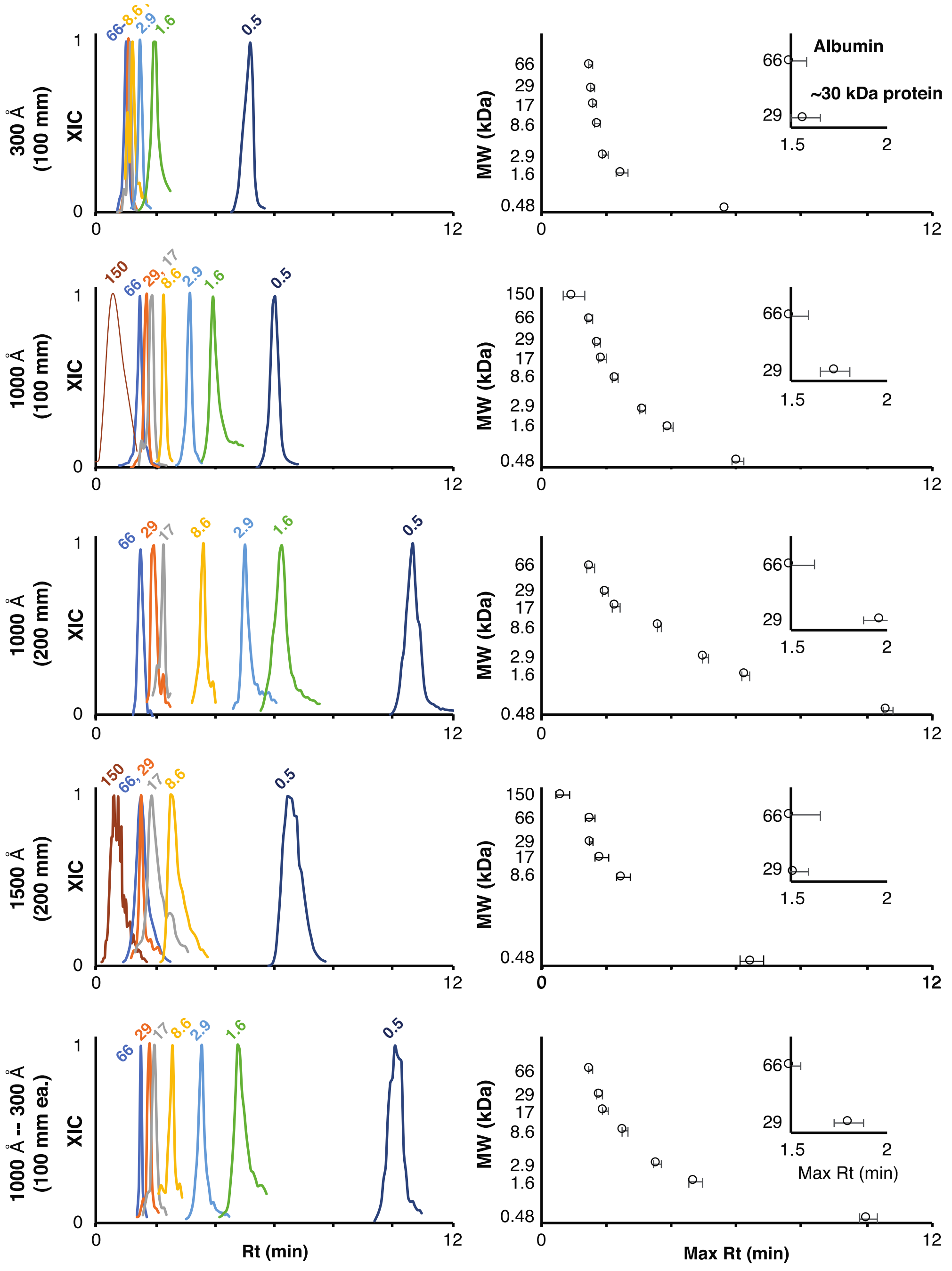

**Figure S2: Performance of different PolyHEA columns.** MS extracted ion chromatogram (XIC) and SEC calibration curves for standard protein/peptides (1 µL injection volume) run on PolyHEA columns ($\varnothing =$ 2.1 mm, 30 µL/min flowrate) of various pore sizes (300, 1000, 1500 Å) and $\ell$ (100, 200 mm). All resin particles were 2 µm except for the 1500 Å particle column (3 µm). The separation of albumin (66 kDa) from proteins < 30 kDa (represented by carbonic anhydrase, 29 kDa) is highlighted (insets). Metrics (Rt, FWHM, albumin-carbonic anhydrase separation) are listed in **Tables S1-2**.

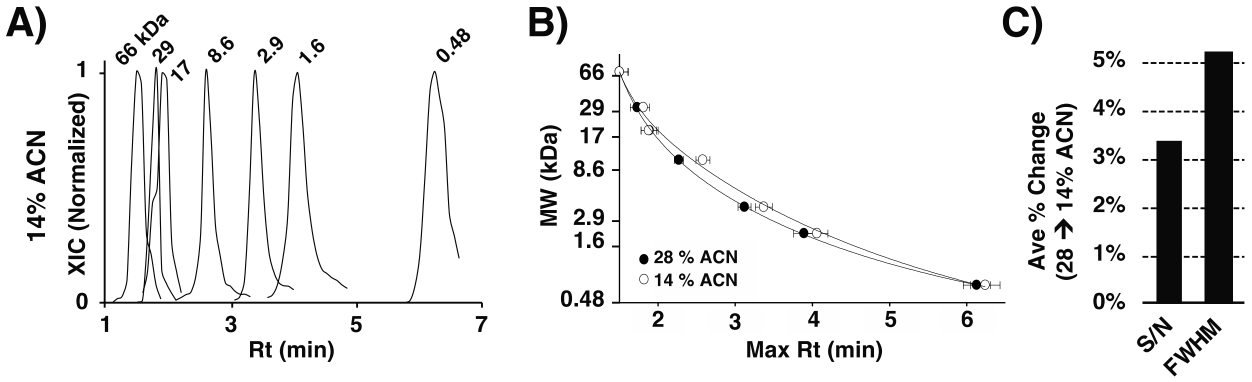


**Figure S3: Comparison of SEC mobile phase organic content.** MS XIC **(A)** and SEC calibration curves **(B)** for seven standard protein/peptides (1 µL injection volume) run on a PolyHEA column ($\varnothing = 2.1$ mm, 1000 Å pore size, $\ell$ = 100 mm, 30 µL/min flowrate) with different mobile phase organic solvent content (14, 28 % acetonitrile, ACN). Calibration curves plot maximum Rt ± half-width half-maximum. Metrics (Rt, FWHM) are listed in **Table S3**. **C)** Chart shows the average percentage change in spectral S/N and SEC FWHM.

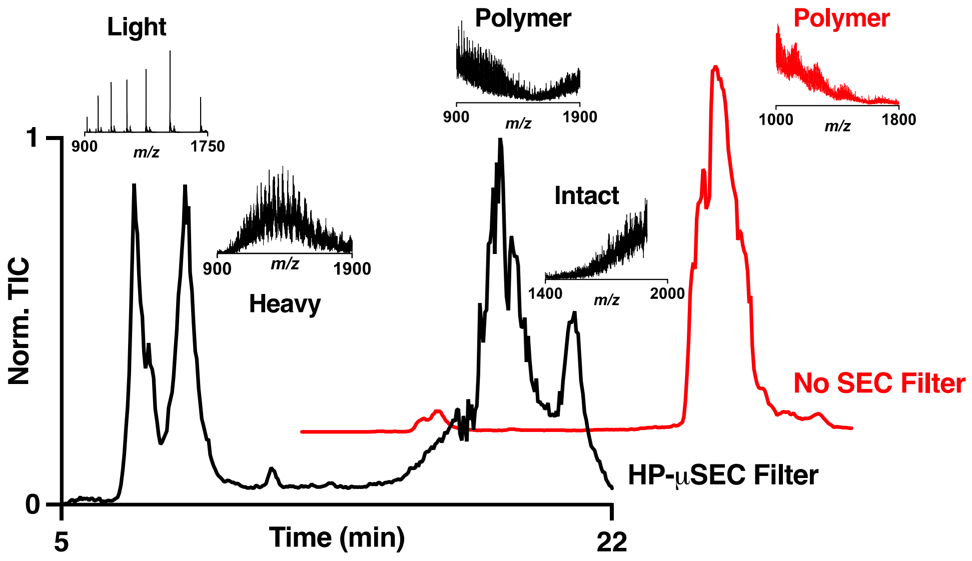


**Figure S4: µSEC-MS platform resolves LMW additives from proteins.** Representative nRPLC-MS chromatograms on a commercial antibody (anti-BNP, isotype IgG) w/o (top, *red*) or w/ (bottom, *black*) online HP-µSEC on a 5 µm particle PolyHEA column (∅ = 1.0 mm, 500 Å pore size, ℓ = 150 mm, 30 µL/min flowrate). Insets show representative mass spectra of peak content. The intact antibody, light chain, and heavy chain were largely overshadowed by background polymer in the No SEC Filter run.

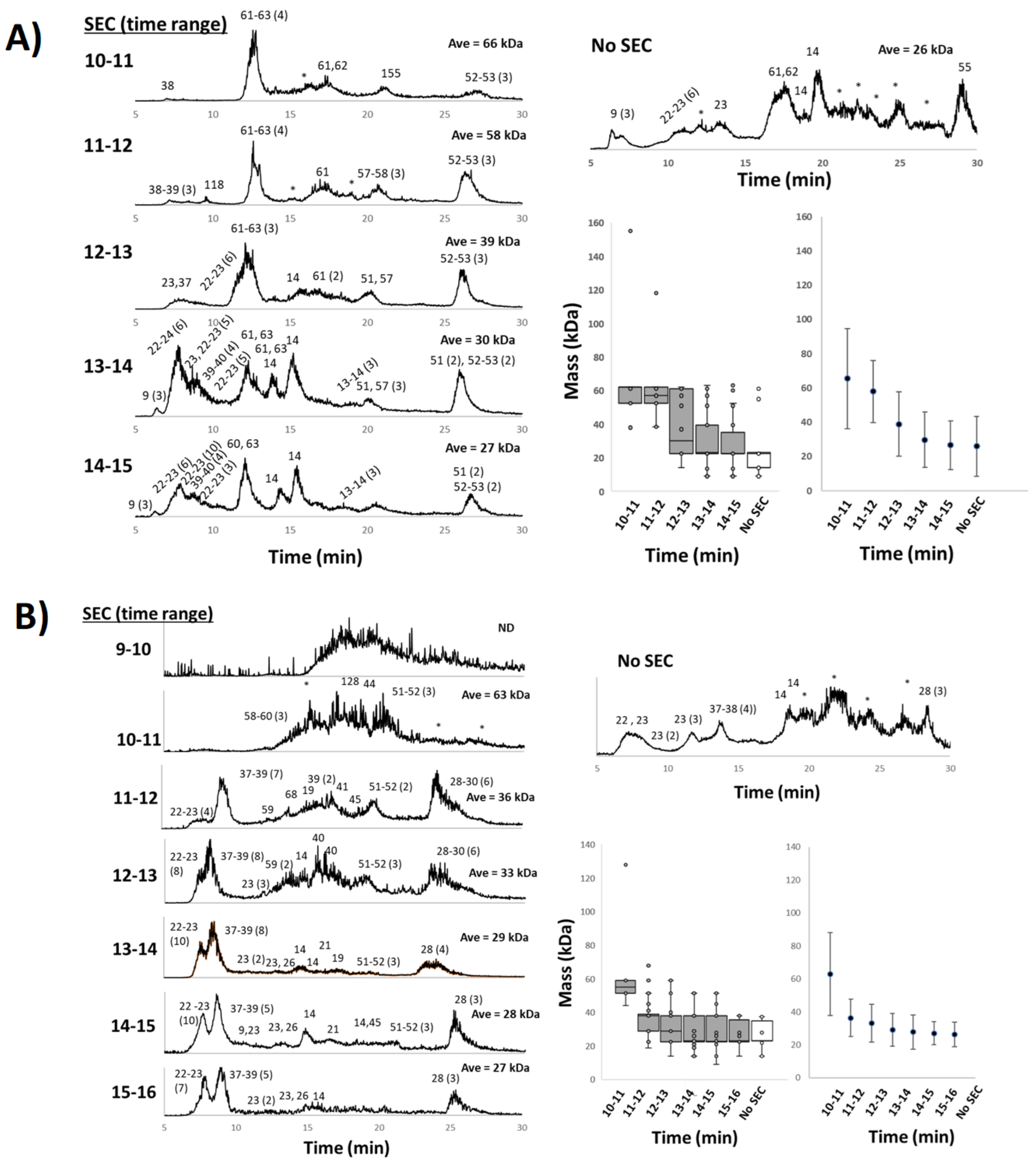


**Figure S5: µSEC-MS platform improves detection of HMW proteins (> 30 kDa) by elimination of LMW additives and proteins (< 30 kDa). (A)** and **(B)** nRPLC-MS chromatograms for various SEC time windows for two NHP CSF IEF fractions (left). Embedded numbers are approximate masses detected across the chromatogram (in kDa). Numbers in parenthesis highlight the number of intact mass tags detected within the respective mass ranges. nRPLC-MS chromatograms for the fractions not subjected to SEC (upper right, **A** and **B**). Box and wisker plots show distribution of masses detected and scatter plots show the estimated average masses observed within each time window (lower right, **A** and **B**). *ND: Areas of chromatograms where *m/z* intensity profile within spectra was suggestive of large protein elution but without resolved charge states.

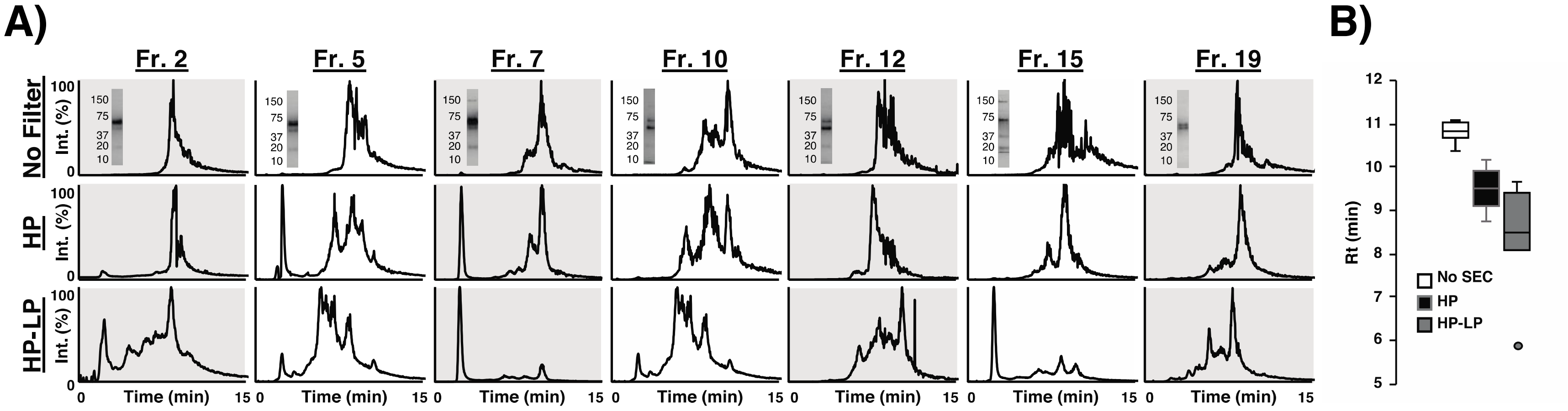


**Figure S6: High- and low-pass filtration increases dynamic range of CSF IMT detection from OFFGEL IEF fractions. A)** LC-MS chromatograms for selected CSF IEF fractions across the pI range (**Figure S6-left**) analyzed w/o SEC ("No SEC", top); w/ high-pass µSEC ("HP", middle); and w/ both high- and low-pass µSEC ("HP-LP", bottom). When no filter is applied, the chromatograms are typically dominated by one peak for an abundant high mass serum-derived proteins (e.g., albumin or transferrin) that elute later in the RPLC gradient. With each added stage of filteration there is a general shift in weighted average total ion signal observed to the polar end of the RPLC gradient **(B)** that corresponds with a 20.5x reduction in S/N and 35.4 % reduction in #s of HMW IMTs (> 30 kDa) and a 13.2x increase in S/N and 167 % increase in #s of LMW IMTs (< 30 kDa) observed (**Figure 5**).

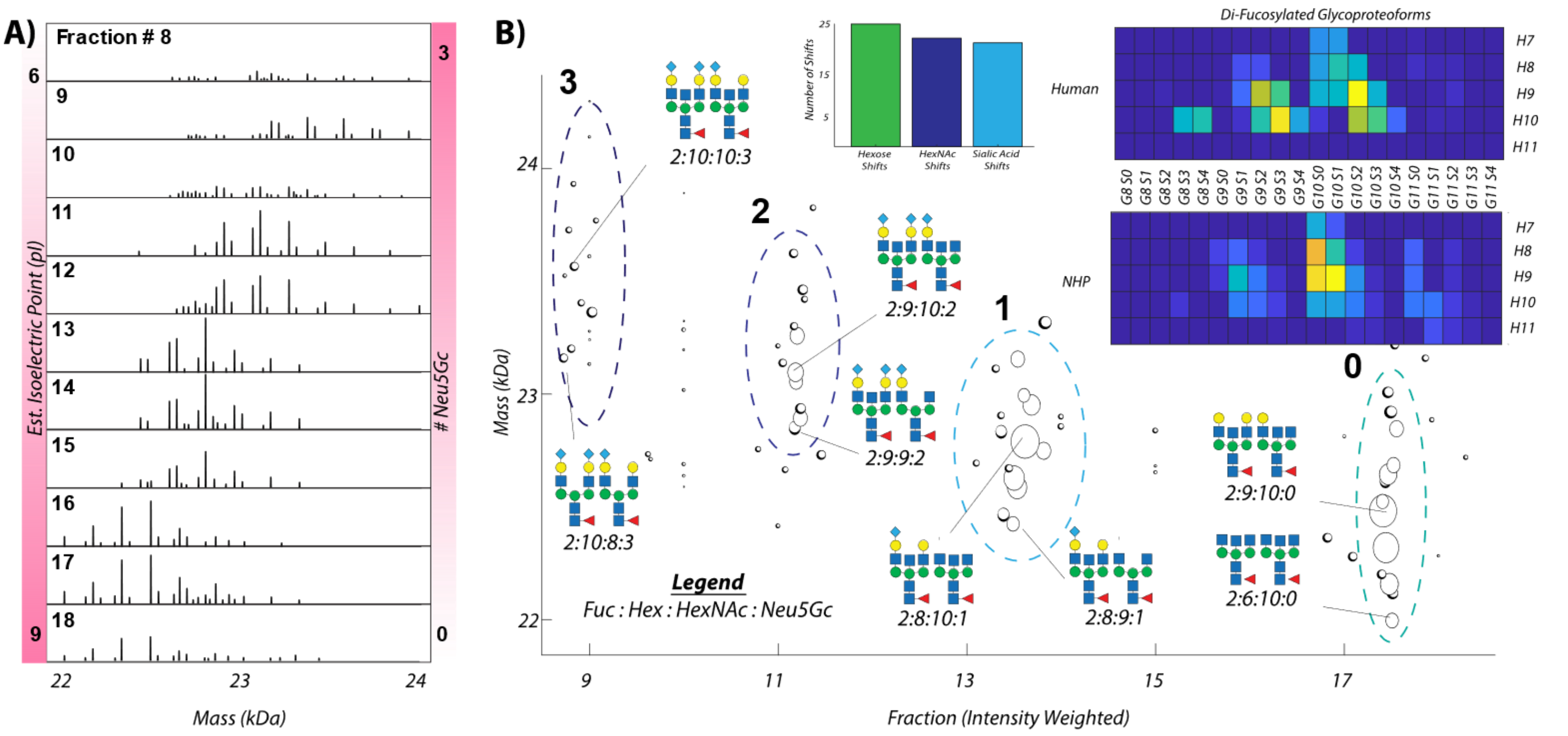


**Figure S7: µSEC²-nRPLC-MS platform enables detection of di-N-glycosylated L-PGDS from OFFGEL IEF fractions obtained on NHP CSF. (A)** Deconvoluted mass spectra associated with L-PGDS glycoforms observed in the respective IEF fractions. **(B)** 2D mass vs. pI plot for binned data for the observed putative di-fucosylated, di-N-glycosylated L-PGDS glycoproteoforms grouped by sialic acid (S) content. Inset bar chart shows frequency of delta mass shifts associated with the highlighted monosaccharides. The data suggests that with decreased pI, the number of siaylic acid residues on each glycoproteoform increases. Inset heatmaps compare the glycoproteoform sugar composition observed for the current NHP data vs. that obtained previously on human CSF L-PGDS,[2] here using ~80-fold less starting material. F = fucose (Fuc); H = hexose (Hex), G = N-acetyl hexosamine (HexNAc). For NHP L-PGDS, S = N-glycolylneuraminic acid (Neu5Gc); for human L-PGDS, S = N-acetylneuraminic acid (Neu5Ac).
